# A hemagglutinin and neuraminidase biased immunological memory shapes the dynamics of antibody responses to Influenza A virus

**DOI:** 10.1101/2024.03.14.584765

**Authors:** Xia Lin, Jiaqi Wang, Shiman Ling, Cheng Xiao, Zaolan Liang, Cheuk Long Chow, Bingyi Yang, Biying Xiao, Benjamin Cowling, Richard Webby, Mark Zanin, Sook-San Wong

**Affiliations:** Guangzhou Medical University, Xinzao, Panyu District, Guangzhou, 511436, P. R. China; State Key Laboratory for Respiratory Diseases, 195 Dongfengxi Rd Guangzhou, 510182, P.R. China; Pasteur Research Pole, School of Public Health, LKS Faculty of Medicine, 5 Sassoon Road, The University of Hong Kong, Hong Kong SAR, China; World Health Organization (WHO) Collaborating Centre for Infectious Disease Epidemiology and Control, School of Public Health, The University of Hong Kong, Hong Kong SAR, China; Department of Infectious Disease, St. Jude Children’s Research Hospital, Memphis, Tennessee, USA; School of Public Health, LKS Faculty of Medicine, The University of Hong Kong, Hong Kong SAR, China; Center for Immunology & Infection Limited, Hong Kong SAR, China

**Keywords:** immunological imprinting, H3N2, hemagglutinin, neuraminidase, antibody landscape

## Abstract

Influenza A virus (IAV) infection establishes a more diverse immunological memory to different viral proteins compared to vaccination. We hypothesized that the relative abundance of pre-existing immune memory to different viral antigens could skew post-infection antibody responses. To explore this, we generated mouse models with either an IAV hemagglutinin (HA)- or neuraminidase (NA)-biased immunological memory. We inoculated groups of mice with cocktails of isogenic viruses bearing antigenically-distinct HA (H3v) or NA (N2v) chosen to span the IAV H3N2 human circulation history. We challenged the mice with two H3N2 strains of opposing virulence and antigenic distance (AD) and examined the post-infection antibody landscapes. In both challenges, immune-naïve mice seroconverted to both HA and NA whereas in primed mice, antibody response was detected to the antigen for which there is no pre-existing memory. In cases where the homologous antibody response was blunted, there was diversification on the breadth of response to antigenically-related strains with low baseline titers. Our findings clarifies the concept of “original antigenic sin” and demonstrate a mechanism by which the dynamics of antibody responses to HA and NA after infection can be altered by pre-existing immunity.

## Introduction

The influence of pre-existing immunity on influenza vaccine and post-infection antibody responses has been well documented in many instances. The first influenza virus (IAV) exposure imprints the immunological memory, resulting in high antibody titers to the first and other antigenically similar strain, that can persist for life. This imprinting can bias subsequent responses to infections or vaccinations, resulting in misdirected antibody responses after vaccination(Corti et al., 2011; Ellebedy et al., 2014; Huang et al., 2015; Lessler et al., 2012; Reber et al., 2015; Ross et al., 2014; Schmidt et al., 2015) or even impacting disease susceptibility(Gostic et al., 2019). In another example, repeated influenza vaccination has also been associated with reduced post-vaccination antibody responses and potentially reduced vaccine effectiveness although the underlying immunological mechanisms of this is unclear(Jones-Gray et al., 2023). Research progress in this area is largely impeded by the difficulties in capturing the complex human immune history to influenza either in human studies or in animal models. Further, most of our current understanding has been based on immunity to the IAV hemagglutinin (HA) protein, the major antigenic target on the virion and immunity to other viral proteins, such as neuraminidase (NA) has been relatively understudied(Krammer et al., 2019).

In a previous study on the burden of community acquired influenza in Auckland, New Zealand, we found that adults and children demonstrated a different post-infection antibody profile to HA and NA. Adults older than 20 years of age demonstrated a preferential induction of antibodies against either HA or NA, while children were more likely to generate antibody responses to both HA and NA(Wong et al., 2020). This suggests that immune history may play a role in determining the dynamics of antibody response to different viral proteins. Such observations of “intravirionic competition” between HA and NA have been reported and elegantly explored in works by Kilbourne et al(Johansson et al., 1987; Kilbourne et al., 1987). Through a series of studies, they showed that i) immune responses to HA and NA are interlinked during natural infections and vaccination ii) that pre-existing HA immunity can reduce antibody response (damping) to NA during subsequent infections if the HA is similar and iii) this process was primarily mediated at the memory B cell level. However, these studies were conducted based on limited strains and did not reflect the H3N2 antigenic evolution in humans to date. To further complicate matters, as influenza vaccines are HA-enriched, regularly vaccinated older adults, such as those in New Zealand (data based on https://stats.oecd.org), may have a more HA-dominant immunological memory compared to unvaccinated populations(Dugan et al., 2020). How this will impact the dynamics of antibody responses to HA and NA is unknown.

To explore how pre-existing immunity, examined in the context of H3N2 antigenic evolution in the last 50 years, can influence the induction of HA or NA antibodies after IAV infection, we generated mouse models with diverse and antigenically-distant HA or NA immune memory using a cocktail of recombinant IAV. Previous studies have attempted sequential infections in animal models, but these experiments were often limited by the number of exposures due to the time needed between infections(Carter et al., 2017; Chiba et al., 2023). Although our approach of cocktail IAV inoculations in mice does not accurately replicate the human sequential exposure, our goal was to reconstitute an immunological memory to diverse antigenic strain that would approximate the exposure history of an older adult; i.e someone who was first exposed to the 1968 H3N2 when it first emerged in the human population. For this reason, we chose strains that are sufficiently antigenically distant (approximately 10 years apart) that could reasonably be expected to re-infect humans during their lifetime. We studied subtype H3N2 IAVs as their antigenic history in humans is well-characterized, allowing the selection of IAVs with sufficient antigenic distance(Bedford et al., 2014; Sandbulte et al., 2011; Smith et al., 2004).

This study had two aims; (i) to understand and disentangle the determinants of antibody dynamics to influenza HA and NA, and (ii) to determine the consequences of a biased immunological memory on antibody responses to IAV HA and NA during subsequent infections. Our findings will have important implication in how we interpret seroepidemiological findings to influenza or other highly antigenically variable pathogens such as SARS-CoV-2.

## Results

### Cocktail priming with diverse HAs induced a balanced antibody response

To establish an HA-biased immune memory, we generated recombinant H3_V_N1 rg viruses bearing H3s that represented distinct HA antigenic clusters(Smith et al., 2004) and the N1 from A/Puerto Rico/8/1934 (H1N1) (PR8). For the period of 1968 to 2019, we selected A/England/42/1972 (H3N2) (EN72), A/Wuhan/359/1995 (H3N2) (WU95), A/Perth/16/2009 (H3N2) (PE09) and A/Hong Kong/4801/2014 (H3N2) (HK14). To ensure comparable replicative fitness of H3_V_N1 viruses, we assessed the growth kinetics of these viruses in MDCK and hMDCK cells, the latter which expresses more α2,6-linked sialic acid, which is preferred by mammalian adapted IAV. H3_EN72_N1 exhibited slower replication kinetics compared to other strains in both cell lines while H3_PE09_N1 exhibited a slight replicative advantage at 24 and 36HPI in MDCK cells compared to hMDCK cells (Supplementary Fig. 1a-b). Overall, although H3_V_N1viruses exhibited some variability in their initial replication rates, all eventually replicated to comparable peak titers within 1 log by 36HPI.

We induced HA-biased immune memory by inoculating mice with a **c**ocktail of diverse **H**A (**CH**); H3_EN72_N1, H3_WU95_N1, H3_PE09_N1 and H3_HK14_N1 (Table 1) either simultaneously or sequentially in two doses. The viruses were inactivated and administered intraperitoneally in equivalent amounts (CH1). To ensure a potent antibody response, we also included an adjuvanted regimen (CH2) and a regimen using live viruses (CH3)(Atmar et al., 2006; Gautam et al., 2020; Gautam et al., 2019). Of note, we also included a **s**equential **H**A priming (**SH**) and **s**equential **H**A and **N**A priming (**SHN**) group. However, we found that this approach could not elicit comparable response to all the antigens; hence we concentrated on responses elicited by the cocktail immunization strategy. Data from sequential priming is provided as supplemental material (Supplemental Fig. 2 and 3) for reference.

**Table 1.** Recombinant viruses encoding H3^a^_V_N1 or H6^b^N2_V_. Unless otherwise indicated, all variable HA or NA (in bold) are derived from H3N2.

| **H3<sub>v</sub>N1<sup>a</sup>** | **HA from strain(Smith et al., 2004)** | **H6<sup>b</sup><sub>N2v</sub>** | **NA from strain(Sandbulte et al., 2011)** |
| --- | --- | --- | --- |
| **H3<sub>EN72</sub>N1** | A/England/42/1972 | **H6N2<sub>EN72</sub>** | A/England/42/1972 |
| **H3<sub>WU95</sub>N1** | A/Wuhan/359/1995 | **H6N2<sub>WU95</sub>** | A/Wuhan/359/1995 |
| **H3<sub>PE09</sub>N1** | A/Perth/16/2009 | **H6N2<sub>PE09</sub>** | A/Perth/16/2009 |
| **H3<sub>HK14</sub>N1** | A/Hong Kong/4801/2014 | **H6N2<sub>HK14</sub>** | A/Hong Kong/4801/2014 |

<sup>a</sup>N1 from A/Puerto Rico/8/1934 (H1N1)
<sup>b</sup>H6 from A/Teal/Hong Kong/W312/1997 (H6N1)

The cocktail immunization elicited HI titers against EN72, WU95, PE09 and HK14 that were boosted comparably in all immunized groups after two doses (Panel CH1 to CH3, Fig. 1b-d). There was a trend of higher titers and lower fold-changes elicited by CH2 compared to CH1, but this difference was not statistically significant. Similarly, no statistically significant differences were evident between CH3, CH2 or CH1 for all antigens. The addition of the adjuvant, Addavax had only minimal effects, since only titers to EN72 were higher after the first dose in CH2 mice compared to CH1 (Fig. 1e-f). Although the NI-antibody titers to the non-variable N1 component also increased after two doses, the average fold-change difference were not statistically significant (Fig. 1g-h).

**Fig. 1.**
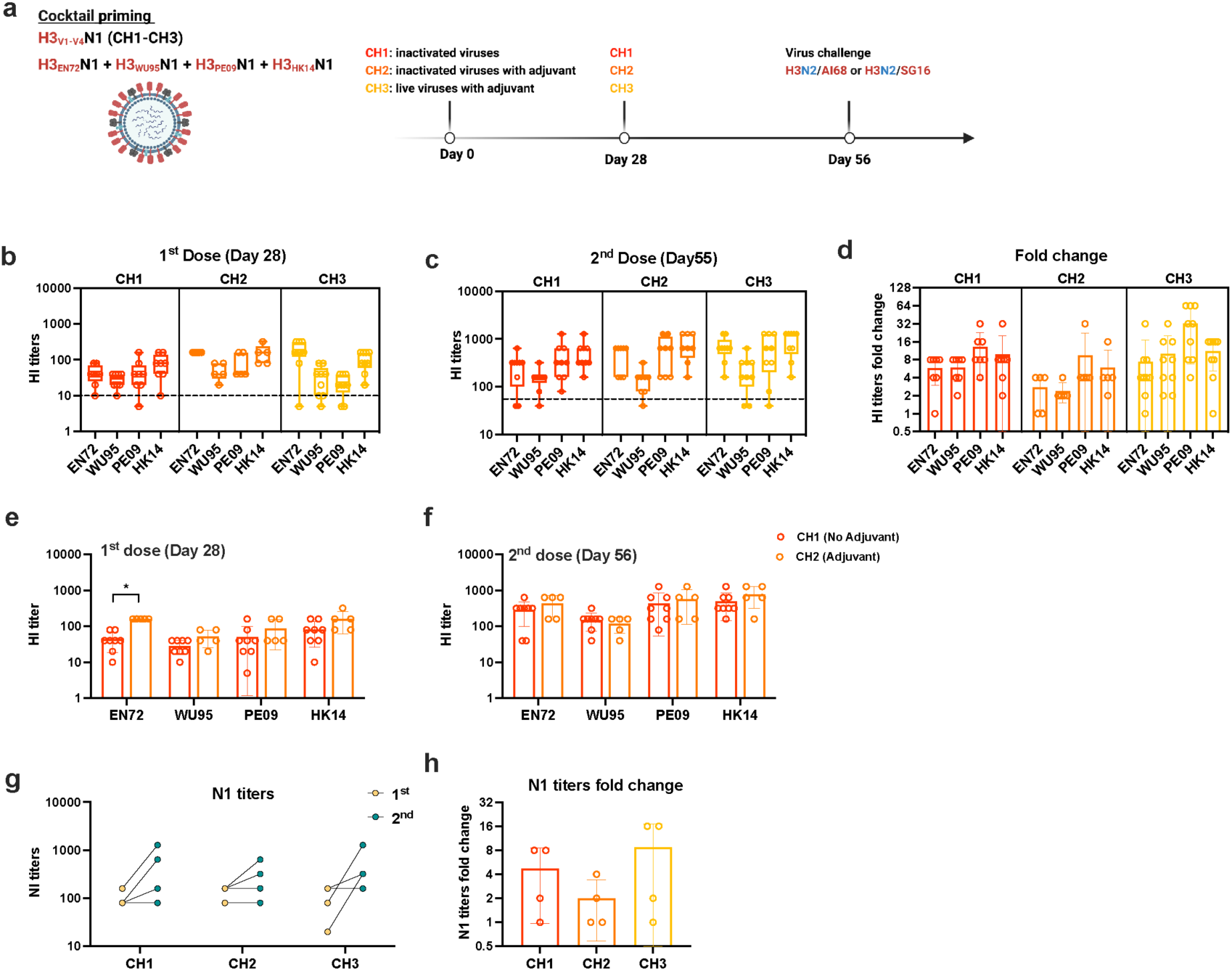
Serum antibody response after cocktail priming with diverse H3. **a** Schematic of HA-imprinted mice. **b-c** Hemagglutination inhibition (HI) serum antibody titers against EN72, WU95, PE09 and HK14 were determined after 1^st^ and 2^nd^ dose. **d** HI antibody fold change between 1^st^ and 2^nd^ dose. **e-f** Comparison of HI titers in mice immunized with adjuvant and without adjuvant after1^st^ and 2^nd^ dose. g**-h** Homologous antigen N1 titers and fold change in H3_V_-imprinted mice. * indicates *P*- value <0.05, ** indicates *P*-value <0.01 and *** indicates *P*-value <0.001 by using unpaired two tailed t-tests. Dotted lines indicated HI titers of 1:10.

When compared to cocktail priming mice, single dose exposure in the SH and SHN groups elicited comparable titers against EN72 and WU95 but not PE09 and HK14, which were significantly lower than that achieved by the cocktail primed mice (CH1-CH3) (*p*<0.001, Supplementary Fig. 2e). Taken together, our results showed that cocktail immunization elicited a balanced antibody response to all antigens while sequential infection with live virus induced a variable antibody response.

### Antibody response occurs only for antigenically related HA with low pre-existing antibody titers

To determine the influence of diverse immunological imprinting on antibody responses after IAV infection, we challenged the cocktail primed mice with two antigenically distant strains. First, we used A/Singapore/INFIMH-16-0019/2016 (H3N2) (SG16), a recently circulating H3N2 strain antigenically closer to HK14 and PE09 than to EN72 and WU95 and is relatively avirulent in mice. At the infecting dose, all mice were asymptomatic (data not shown). Infection of the PBS group elicited modest and comparable HI titers to SG16, HK14 and SW19 (black line in Fig. 2), indicating that these viruses were antigenically similar and HI-responses were of limited breadth. For the primed mice, pre-challenge antibody titers to immunizing strains (from Fig. 1) as well as additional strains that were tested to validate the shape of the landscape, were plotted as bars in Fig. 2. As expected, post-infection antibody landscapes were considerably broader in primed mice, with a largely similar shaped response amongst the different cocktail primed groups. Mice in CH2 and CH3 retained higher HI titers to AI68, EN72 and VI75 strains, respectively, suggesting some immunogenicity advantage in CH2 and CH3 compared to CH1 (Fig. 2a-d). Antibody titers of mice in the cocktail primed groups did not increase to SG16 or HK14 but did increase to past strains of BE89, WI05 and SW13. Pre-challenge HI titers to SG16 were high (GMT 470.3, 95% CI: 294-752.4), while low to BE89 (GMT 5, 95% CI: 5-5), WI05 (GMT 5, 95% CI: 5-5) and SW13 (GMT 17.1, 95% CI: 9.6-30.7) in these mice.

**Fig. 2.**
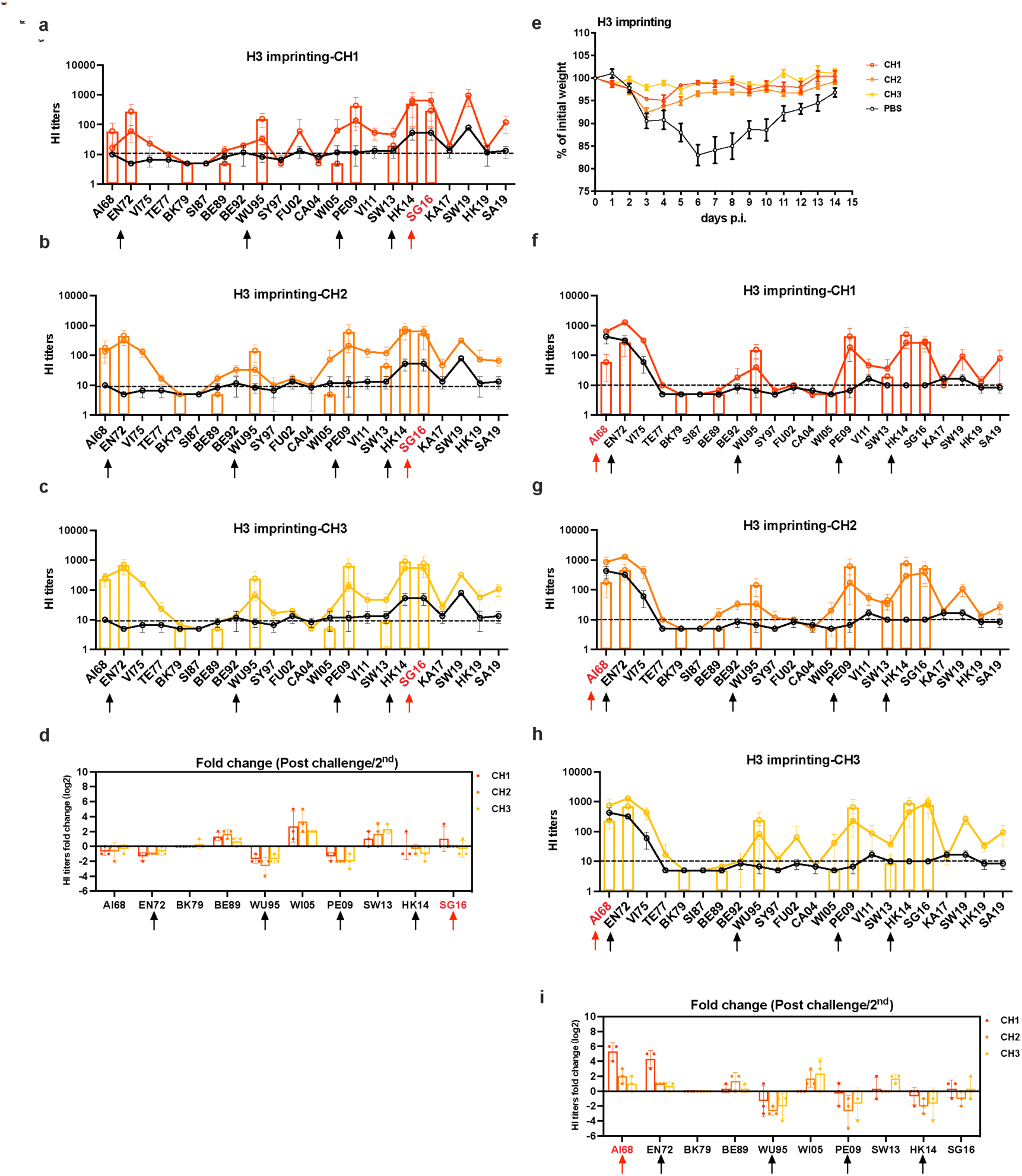
HI antibody landscape after challenge with SG16 or AI68. **a-c** HI titers (solid lines) against 22 historical H3N2 viruses isolated between 1968 to 2019 in cocktail primed H3 mice after infected with SG16. **d** HI titers fold change between pre- (after 2^nd^ dose or infection) and post challenge with SG16 in all groups. **e** 21 days after final vaccination, H3_V_-imprinted mice were infected with 10^4^ PFU AI68 and weight loss were recorded daily for 14 days. **f-h** HI antibody landscape against 22 historical H3N2 viruses in cocktail primed mice infected with AI68. **i** HI titers fold change between post challenge with AI68 and 2^nd^ dose vaccination. The histogram indicated the pre-challenge antibody titers to immunizing antigens (black arrows) and selected strains that were chosen to verify the shape of the pre-challenge landscape curve. The black line indicated titers in the PBS group challenged with SG16 or AI68. The red arrow indicated challenge virus. Dotted lines indicated the 1:10 in HI titers.

**Fig. 3.**
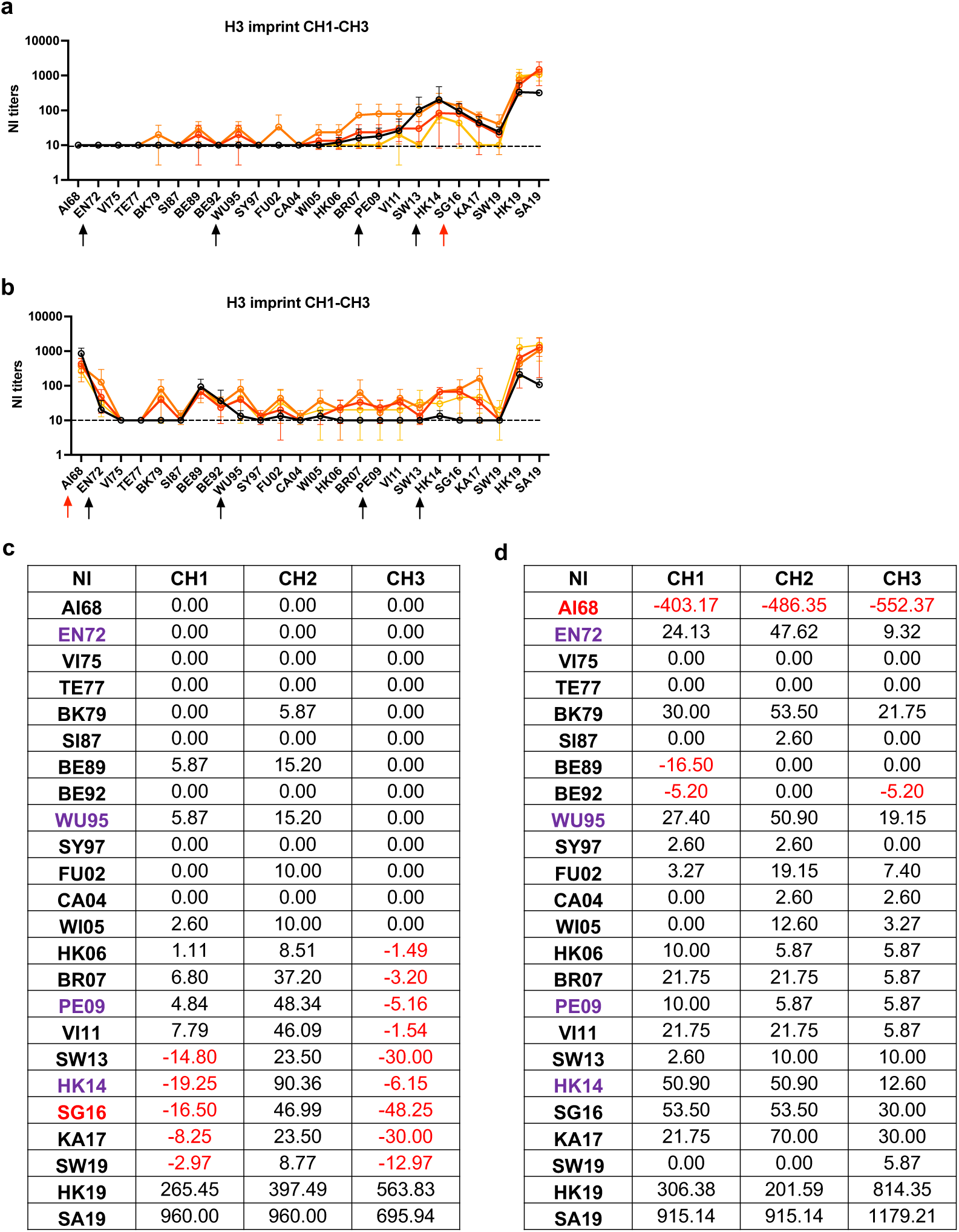
NI antibody response in H3_V_-imprinted mice. **a-b** NI antibody response against 24 historical H3N2 viruses isolated between 1968 to 2019 in H3_V_-imprinted mice after challenged with SG16 and AI68. **c** Antibody response after subtracted the NI geometric mean titer (GMT) response for each strain in each immunization group from PBS. Negative values were highlighted in red. Dotted lines indicated the 1:10 in NI titers. The challenged strain was highlight in red and priming strains were in purple.

We also challenged the mice with an older H3N2 strain, A/Aichi/2/1968 (H3N2) (AI68), which is antigenically closer to EN72 than WU95, PE09 and HK14 and is highly pathogenic in mice. The challenge dose elicited a 15% weight loss in the PBS group by 6DPI, while other groups lost less or no weight (Fig. 2e). Mice inoculated with live viruses (CH3) showed no significant weight loss compared to those that received inactivated antigens CH1 and CH2, which lost 5 to 7% starting body weight by 3DPI.

Infection of the PBS group with AI68 elicited HI titers to AI68, EN72 and VI75. Titers to these three strains were also higher in the immune mice (Fig. 2f-h) and were boosted the most compared to other tested strains (Fig. 2i). CH1, which had the lowest pre-challenge titers, showed the largest antibody increase to AI68 and EN72 while minimal increases were observed for CH2 and CH3.

Taken together, after infection with SG16 or AI68, the magnitude of the antibody response elicited depends on pre-existing titers of even the antigenically-related antigens and the breadth of response varied according to the degree of pre-existing immunity established (Fig. 2). Minimal cross-reactivity is observed to antigens that are sufficiently distant from infecting strains.

### NA antibody landscapes after challenge with antigenic distinct H3N2

To explore how the pre-existing immunological memory to diverse HA influences the NA antibody landscape, we measured NI titers against a panel of corresponding H3N2 IAVs after challenge SG16 or AI68 (Fig. 3a and b). As relatively large amounts of sera are required to perform ELLAs, NI antibody landscapes were only constructed based on post-challenge sera.

Despite the lack of symptoms and an antibody response to HA after challenge with SG16, seroconversion (four-fold increase in antibody titer) was still detected against N2 in CH1 and CH2, suggesting that the high HI titers to SG16 did not provide sterilizing immunity. As observed with HA, antibodies were largely boosted to closely related antigens in influenza-naïve and cocktail immunized mice, although titers to HK19 and SA19 were also unexpectedly high. The AI68-infection also boosted antibody titers to N2 of BE89 and BE92, highlighting a possible antigenic cross-reactivity between the N2 of AI68 with BE89 and BE92. In addition, compared to the PBS group, CH1, CH2 and CH3 also showed responses to antigenically distant strains after both challenge, suggesting that pre-existing antibodies to heterosubtypic NA (N1 in this case) can help boost the breadth of N2 antibody responses.

To evaluate if there is quantitative difference in NI response between primed and naïve mice, we subtracted the NI GMT response for each strain in each immunization group from PBS (Fig. 3c-d). Negative values indicate responses that are less than naïve mice, suggesting a potential “damping” effect from prior immunity(Johansson et al., 1987; Kilbourne et al., 1987). Quantitatively, CH1 and CH3 demonstrated less NI antibodies against SG16 and its closely related strains, SW13, HK14, KA17 and SW19, while CH2 showed comparable responses to the PBS-immunized group. This difference was not due to pre-challenge N1 antibody profiles as titers were comparable between CH1, CH2 and CH3 (Fig. 1h), suggesting that the NA antibody response is “dampened” in the presence of prior immunity to HA or an irrelevant NA and this effect was alleviated with the use of adjuvant in CH2.

### Antibody profile of diverse NAv-imprinted mice

To study the effect of immunological memory to diverse NAs, we generated recombinant H6N2_V_ IAVs bearing antigenically distinct N2s from H3N2(Sandbulte et al., 2011) IAVs in the backbone of a constant H6. The growth kinetics data showed that titers of H6N2_V_ IAVs plateaued after 24HPI and 48HPI in HMDCK cells and MDCK cells, respectively (Supplemental Fig. 1c-d). Unlike the H3vN1 IAVs, there were no significant differences between the viral titers of different H6N2_V_ IAVs at any time point. Therefore, the different N2s had minimal effects on replication kinetics.

Using the same immunization regimen as before, two doses of cocktail priming induced comparable NI titers to most of the antigens in CN1, CN2 and CN3 (Fig. 4b-d), except to WU95, which unexpectedly, was not boosted by the second dose. Higher NI titers against EN72 were observed in CN2 compared to CN1 after the first and second dose, thus indicating an adjuvant effect (Fig. 4e-f). Responses to the homologous antigen H6 were also boosted after second dose vaccination N2_V_-imprinted mice (Fig. 4g-h).

**Fig. 4.**
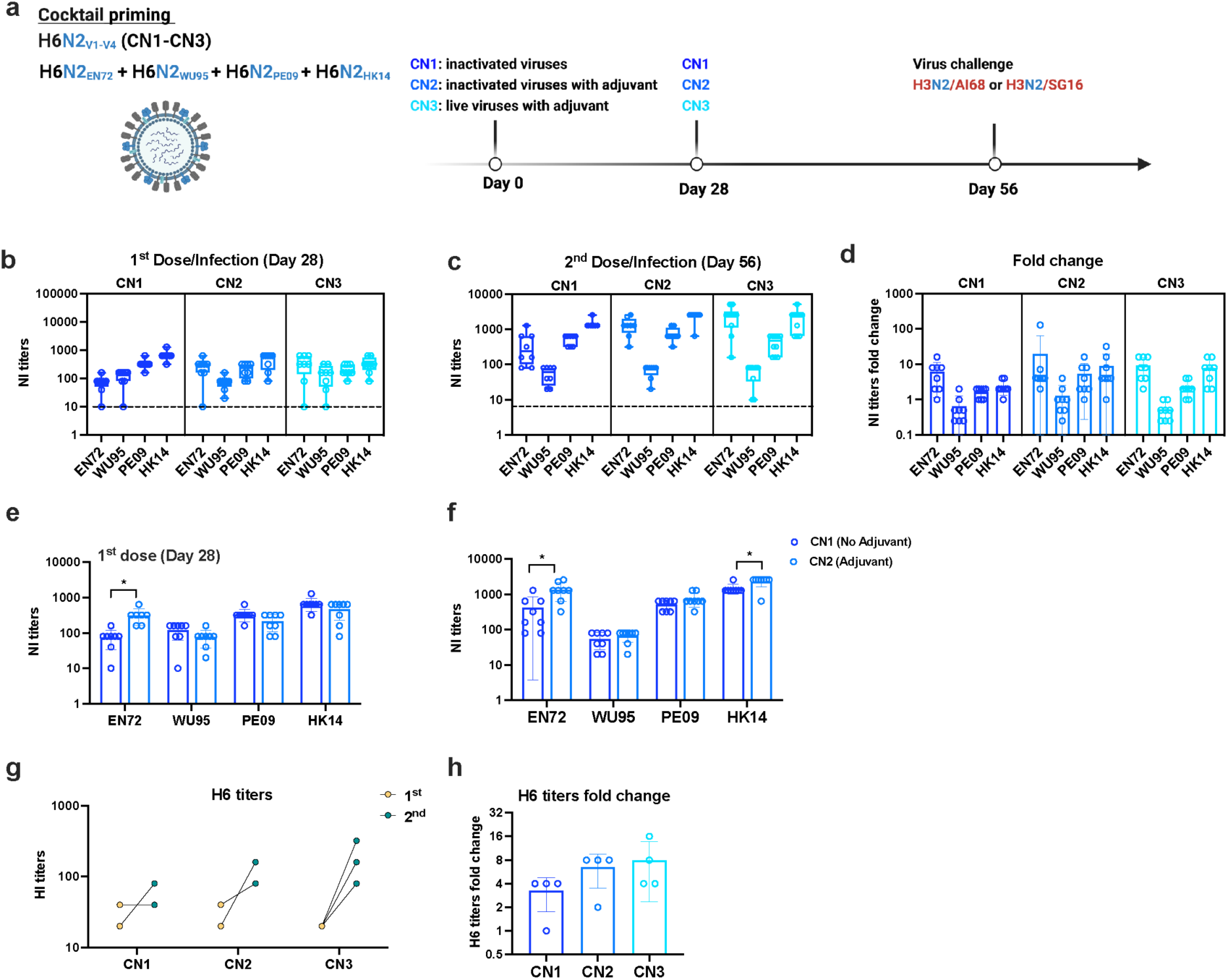
Serum antibody response after cocktail priming with diverse N2. **a** Schematic of NA-imprinted mice. **b-c** Neuraminidase inhibition (NI) sera antibody titers against EN72, WU95, PE09 and HK14 were evaluated via ELLA assay for each group after 1^st^ and 2^nd^ dose. **d** NI antibody fold change between 1^st^ and 2^nd^ dose. **e-f** Comparison of NI titers in mice immunized with adjuvant and without adjuvant after1^st^ and 2^nd^ dose. **g-h** Homologous antigen H6 titers and fold change in N2_V_-imprinted mice. * indicates *P*-value <0.05, ** indicates *P*-value <0.01 and *** indicates *P*-value <0.001 by using unpaired two tailed t-tests. Dotted lines indicated the 1:10 in NI titers.

### NAv-imprinted mice elicited broader NA antibody response after infection with antigenic distinct H3N2

After challenge with SG16, mice in the PBS group only elicited NI titers to strains in circulation after PE09 and to HK19 and SA19, consistent with data seen in Figure 3a. In contrast, cocktail primed mice elicited NI antibodies against most H3N2 IAVs in the panel except for strains circulating prior to WU95 (Fig. 5). NA reactivity in CN1, CN2 and CN3 was highest to HK14 NA and NI titers of >1:40 to strains circulating after WU95 were detected (Fig. 5a-c). Although NI titers did not increase to SG16 after infection in all immunized groups, NI antibody titers to WU95, WI05, PE09, SW13 and HK14 were boosted significantly (Fig. 5d).

**Fig. 5.**
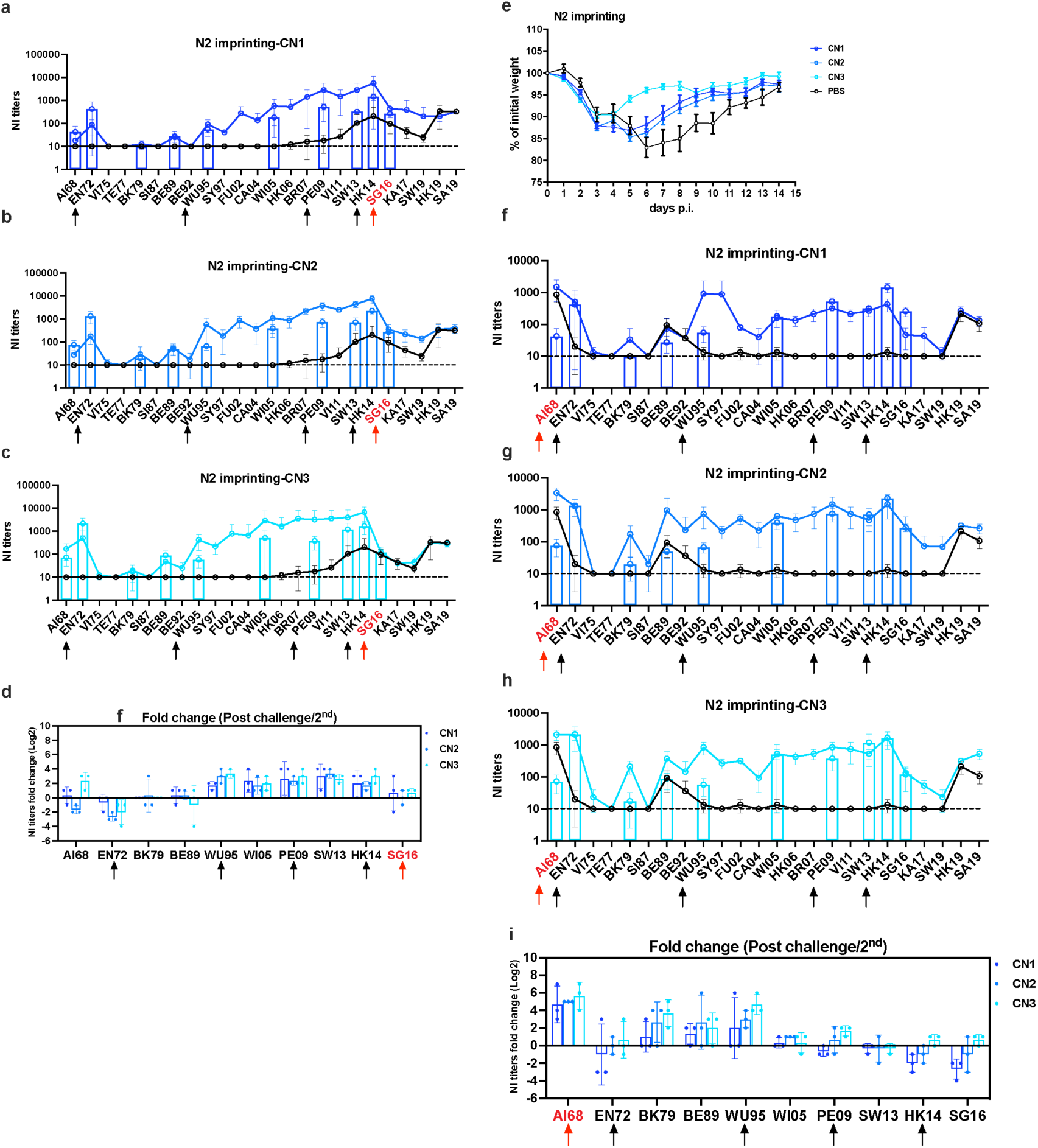
NI antibody landscape in N2_V_-imprinted mice after challenged with SG16 or AI68. **a-c** NI titers were determined by ELLA assay against 24 historical H3N2 viruses in cocktail priming N2 mice after infected with SG16. **d** NI titers fold change between post challenge with SG16 and 2^nd^ dose. **e** 21 days after final vaccination, N2_V_-imprinted mice were infected with 10^4^ PFU AI68 and weight loss were recorded daily for 14 days. **f-h** NI antibody landscape against 24 historical H3N2 viruses in cocktail priming N2 mice infected with AI68. **i** NI titers fold change between post challenge with AI68 and 2^nd^ dose vaccination. The histogram indicated the pre-challenge antibody titers to immunizing antigens (black arrows) and selected strains that were chosen to verify the shape of the pre-challenge landscape curve. The black line indicated titers in the PBS group challenged with SG16 or AI68. The red arrow indicated challenged virus. Dotted lines indicated the 1:10 in NI titers.

Diverse NA imprinted mice were also inoculated with AI68 (Fig. 5e). All mice lost comparable weight within the first 3DPI, but CN3 mice regained their weight quicker than CN1 and CN2 mice. PBS mice recovered the slowest, taking 14 days to recover to 95% of their original weight, while primed mice only took 9 days. These weight loss curve showed that NA immunity provided only partial protection against AI68, in agreement to published data(Strohmeier et al., 2021). After challenge, NI antibodies fold changes were highest against AI68 in CN1, CN2 and CN3, and with the exception of EN72, NI-antibody fold changes were progressively less up to WU95 (Fig. 5f-i).

Fig. 6 shows the post-infection HI antibody landscapes in the diverse NA-imprinted mice. As before, although NI seroconversion to SG16 were not observed in the immunized mice, HI titers were elicited against H3, indicating successful infections. However, unlike the results described in Fig. 4, antibody responses to HA were more strain specific compared to NA. Unlike NA, pre-existing antibodies to H6 did not impact the breadth of post-challenge antibody response to H3 as immunized groups displayed similar HA cross-reactivities compared to influenza-naïve groups. All the groups showed narrow responses, recognizing only HK14, SG16 and SW19 after challenge with SG16 and recognizing only AI68, EN72 and VI75 after challenge with AI68. Following subtraction of responses detected in PBS, lower HI titers to SG16, AI68 and related strains were detected in CN1 and CN3. This difference is not likely due to severity of infection as morbidity was similar in CN1 and CN2. As above, this data suggests that Addavax likely increased the magnitude of the post-infection antibody response.

**Fig. 6.**
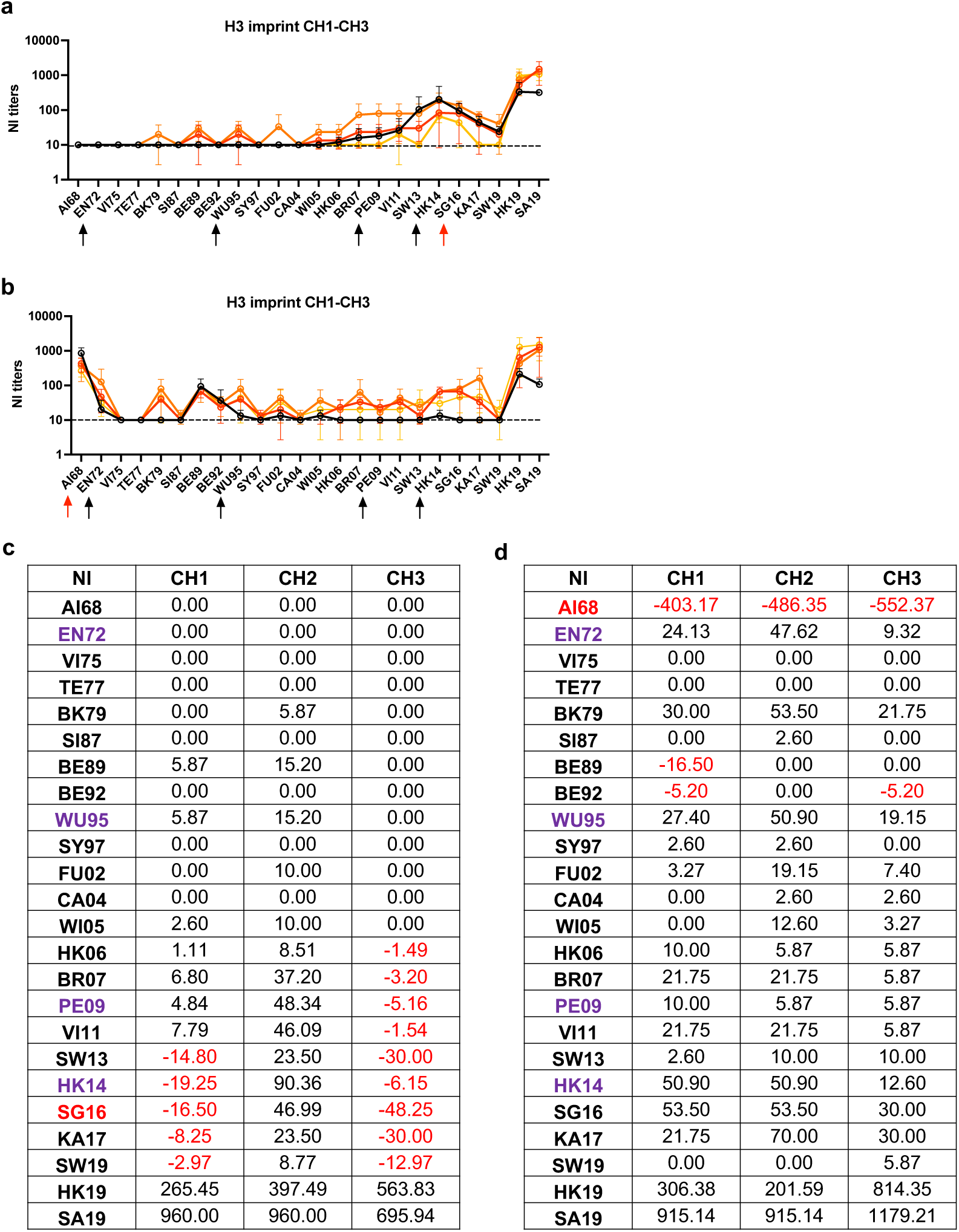
HI antibody response in N2_V_-imprinted mice. **a-b** HI antibody landscape against 22 historical H3N2 viruses in N2_V_-imprinted mice after challenged with SG16 and AI68. **c** Antibody response after subtracted the HI GMT response for each strain in each immunization group from PBS. Negative values were highlighted in red. Dotted lines indicated the 1:10 in HI titers. The challenged strain was highlight in red and priming strains were in purple.

### Antigenic distance and antibody response in HA or NA imprinted mice

We next asked if the antibody responses could be explained within the context of antigenic distance. We estimated antigenic distance (AD) amongst the H3 or N2 proteins using the sequence-based method, based on mismatches at identified antigenic sites(Anderson et al., 2018) (Supplemental Table 2 and 3). AD is calculated with reference to the infecting strain, which was set to “0” in Fig. 7. With this method, the AD between the HA of AI68 to SG16 is 9.02, while for NA is 5.49 (Fig. 7a and d). Using this metric, we noted that our four chosen antigens were not equally spaced in terms of AD. The AD was greatest between EN72 and WU95 (6.12 for HA and 2.75 for NA), whilst they were smallest between PE09 and HK14 (0.84 for HA and 0 for NA). The AD for HA and NA between the challenged strains and the nearest immunizing antigen were 0.57 and 0.39 respectively for SG16 and HK14, and 1.2 and 1.18 respectively for AI68 and EN72.

**Fig. 7.**
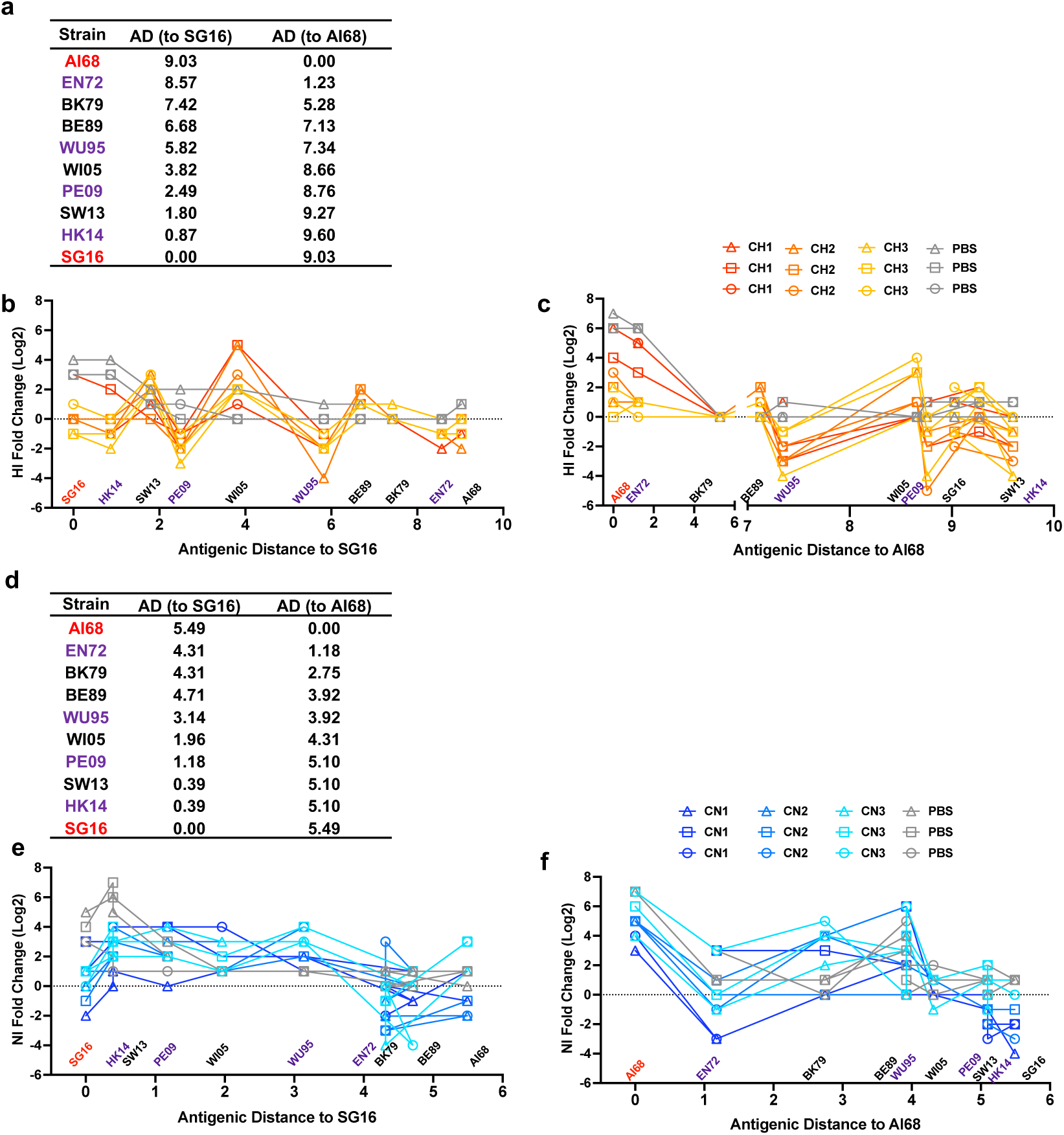
Antigenic distance and antibody response in H3_V_- or N2_V_-imprinted mice. The antigenic distances (AD) for HA and NA between the challenged and the tested strains were calculated by the number of mismatched residues at each antigenic site. **a** Summary of AD to SG16 and AI68 in HA proteins. **b-c** The relationship between antigenic distances and HI antibody fold change (post challenge with AI68 or SG16/pre-challenged titers) in H3_V_-imprinted mice. **d** Summary of AD to SG16 and AI68 in NA proteins. **e-f** The relationship between antigenic distances and NI antibody fold change (post challenge with AI68 or SG16/pre-challenged titers) in N2_V_-imprinted mice. The challenged virus was highlight in red and priming viruses were in purple.

For HA, all mice in the PBS/SG16 challenge group had antibody increases up to SW13 (<2 AD) although the highest responding mice had low level antibodies up to BE89 (∼7 AD) (Fig. 7b). Responses in the PBS/AI68 challenge were uniformly limited to AI68 and EN72 (Fig. 7c). In both challenges however, the naïve mice showed a continuous, even profile across the AD, while responses in the primed mice were punctuated by sharp ‘peaks’ and ‘valleys’. The ‘valleys’ correspond to the point at which the immunizing antigens are located while the ‘peaks’ correspond to the next closest antigen. Notably, the height of these backboosted peaks were inverse of the responses to the challenge strains; i.e the more blunted the homologous response, the larger the backboosted response. The responses in the NA profile within the context of AD were less clear, suggesting this sequence-based AD estimation maybe less accurate for NA.

## Discussion

Studying the how previous exposures influences recall responses in an epidemiologically relevant context is challenging due to the complicated and ‘noisy’ immunological histories in humans that are difficult to replicate in animal models. Contrary to existing strategies, we primed using only diverse H3s or N2s to elicit a broad antibody response and we used viruses rather than proteins or peptides to better reflect intravirionic competition as described previously(Johansson et al., 1987). Through our model, we made several key observations that reflected epidemiological phenomena observed in humans, i) seroconversion to a singular viral protein, for which there is low pre-existing immunity, occurs when there is relatively high preexisting immunity to the alternate protein, ii) infection can occur (as evidenced by seroconversion to NA in SG16 challenge experiment) despite high preexisting HI antibodies and the absence of symptomatic infections and finally, iii) antibody response occurs predominantly only for antigenically related strains for which there is low pre-existing titers and not necessarily in the order of exposure. The last point is an important clarification to the concept of “original antigenic sin”(Francis, 1960); as we demonstrate that only antigenically-related strains are boosted. As such, boosting of antibodies first-experienced in life maybe more relevant in younger individuals.

Further, our data provided a probable cause on why we observed balanced antibody response to HA and NA in children but not adults. The HA antibody landscapes after SG16 and AI68 in both CH and SH mice were similar, suggesting that backboosting will only occur for antigenically-related strains. Data from our sequential priming experiment suggest that the order of exposure was not important, as backboosting was not observed for EN72 or WU95 in the SG16 challenge nor WU95 in the AI68 challenge, which were the first primed strains (Supplemental Figure 3). Although the order of exposure may impact titers of subsequent infections, we show that this “interference” or competition will be less important when the strains are sufficiently distant, consistent with currently proposed models(Miller et al., 2013; Zarnitsyna et al., 2015). Finally, mice imprinted with diverse H3 or N2 antigens elicited less NA or HA antibodies, respectively, to the infecting strains compared to immunologically naïve mice, which is consistent with the works by Kilbourne et al.(Kilbourne et al., 1987). Taken together, our mouse model recapitulated serological phenomena previously reported in humans and showed that the “antibody ceiling effect” is limited to the specific target, and does not reflect an absent immunological response, as antibodies are made towards other related antigens or protein targets. Previous studies by Angeletti et al. had demonstrated that high titers of pre-existing antibodies limited the development of cognate antibody response, driving the diversification of alternative antibody targets that are subdominant in a H1N1-infected mouse model(Angeletti et al., 2017). Studies with SARS-CoV-2 suggests that the circulating antibodies to bind and mask the immunodominant epitopes, resulting in selection and activation of lower affinity B cell clones targeting sub-dominant epitopes at the germinal center(McNamara et al., 2020; Schaefer-Babajew et al., 2023). Our findings were consistent with this phenomenon and further showed that this diversification can result in boosting of HAI-responses to virus strains isolated more than 10 years apart. Thus, some future issues to address would be to define the threshold of this ceiling as well as understand the underlying immunological bottleneck.

We found that, in contrast to sequential infections, cocktail priming elicited comparable antibody responses to all antigens, establishing a “cleaner” system to study downstream antibody responses. H3_V_N1 IAVs showed greater variability in replication kinetics, which we attributed to differences in HA binding preferences, since EN72, isolated only four years after the emergence of H3N2 in 1968, was likely less mammalian adapted compared to the later strains. In contrast, H6N2_v_ IAVs showed comparable replication kinetics in MDCK and hMDCK cells, indicating that HA, rather than NA, was more important for viral replication in these *in vitro* models. One exception was H3_PE09_N1, which failed to elicit significant antibody titers in the sequential infection models despite comparable virus fitness *in vitro*. Contemporary human H3N2 IAVs such as PE09 and SG16, preferred α2,6-linked sialic acid moieties and replicated poorly in the airways of mice which predominantly expresses α2,3-linked sialic acid moieties preferred by avian influenza viruses (such as those bearing the avian-like HA of AI68 and EN72 or the avian H6). Sequential IN inoculation to generate complex immune histories in mice using different IAVs may be influenced by the binding preferences of the viruses. This was not evident when inocula were administered IP, which elicited comparable antibody responses to all antigens, as the peritoneal cavity of the BALB/c mouse expresses both α2,3- and α2,6-linked sialic acids, thus supporting the replication of IAVs with different binding preferences(Gautam et al., 2019; Qi et al., 2009). Although not immediately apparent when comparing the magnitude of vaccine-specific responses, the benefits of the Addavax adjuvant was revealed with the expanded breadth of response as measured by the antibody landscapes, consistent with its reported mode-of-action(Ellebedy et al., 2011).

We selected an antigenically distinct H3N2 IAVs from the past, AI68 and an antigenically distinct contemporary strain, SG16 to investigate the impact of pre-existing H3 or N2 antibodies on the magnitude and breath of antibody response after challenge. AI68 was highly virulent while SG16 was not, partly due to differences as described above. All primed mice were protected from AI68 IAV challenge but the H3 primed mice were better protected compared to the N2-primed mice. This is consistent with the mode of protection mediated by HA and NA antibodies(Allen et al., 2019; Malik and Zhou, 2020; Zanin et al., 2015). Notably, CH3 and CN3, which received IP administration of live viruses were not as well protected against challenge compared to groups that received IN inoculation. This may have been due to poor induction of cellular immunity in the airways of CH3 and CN3 mice(Gautam et al., 2019).

The cocktail immunization strategy with HAs and NAs from antigenically distinct clusters(Sandbulte et al., 2011; Smith et al., 2004) elicited antibodies that could recognize most human H3N2 IAVs, with N2 eliciting responses of greater breadth, spanning strains across a broader time period, compared to H3. This result may have been influenced by the differences in assay sensitivity as well as the slower rate evolution for human N2 compared to H3(Abed et al., 2002; Naeem et al., 2020). Furthermore, the shape of the HI and NI antibody landscapes in the H3_V_-immunized mice were similar to those observed in older adults that is imprinted by the 1968 H3N2 virus, while the response in the naïve mice can represent the immune history of a child(Auladell et al., 2022; Ertesvag et al., 2022; Fonville et al., 2016). The post-infection antibody response in naïve versus immunized mice recapitulated the profile we previously observed in humans; i.e a balanced HA and NA response in naïve mice but a biased HA and NA antibody response in mice with complex influenza immune history. This has important implications in our interpretation of studies relying on singular serological endpoints or our understanding of the correlate of protection for influenza. Assessing a single serological target maybe adequate in a largely naïve or a population with uncomplicated immune history to influenza but maybe insufficient in populations with complex immune history such as older adults or a highly vaccinated population. Further, since our finding suggest that antibodies targeting HA and NA maybe induced differently depending on the exposure history, it is conceivable that this may also impact the mechanisms of protection in different populations, such as that in older adults or those that are repeatedly vaccinated. Finally, we used a sequence-based antigenic distance (AD) approach to quantify the antigenic relationship between various H3 or N2 proteins, which had been developed by Anderson et al. and represented as antigenic cartography of H1N1 viruses(Anderson et al., 2018). This method, as a potential alternative approach to traditional antigenic cartography (HI-based ferret antisera assay)(Smith et al., 2004), can estimate the antigenic relationship across diverse influenza strains. The AD results also supported that HA- or NA-biased mice developed broader antibody response against antigenically-related (quantified by AD) HA or NA, which indicated the dynamics and cross-reactivity of antibody responses to Influenza A virus were based on antigenic distance(Gog and Grenfell, 2002).Unlike HA, NA antigenic sites are less well-characterized, potentially leading to less defined relationship between antigenicity and AD.

Our study has several limitations. The mouse model is limiting in terms of the amount of serum that can be collected, which is particularly relevant in the context of constructing antibody landscapes. Hence, we were only able to measure pre-challenge antibody titers to select strains. Although the distinct antigenicity of the H3 and N2 strains tested here were consistent with data obtained from human sera, there may be distinct species-specific differences(Mestas and Hughes, 2004), particularly in terms of epitope recognition, as was previously shown for H1N1 viruses(Liu et al., 2018). Since the size of the naïve B cells populations and the magnitude of antibody titers measured is smaller in mice compared to humans, it is unknown how this will impact the breadth of antibody responses beyond what is measured in this study(Johnson et al., 2020; Sigal et al., 1987). Finally, our observations are derived from a non-aged mouse model, and as such we are not able to account for differences associated with aging or other conditions that may impact the immune system in older adults.

The landscape generated with our mouse model surprisingly were able to recapitulate the landscapes observed in older adults after infection with HK14 (manuscript in preparation, data available for review purposes). Through this, our findings highlight the importance of baseline antibody titers, which is a proxy for pre-existing immunity, in influencing the dynamics of antibody responses to different viral antigens. This observation may also apply to COVID-19, where most vaccines target the Spike protein and vast majority of the population would have been repeatedly vaccinated. Importantly, infection can occur, despite evidence of high protective antibodies and lack of symptoms. In humans, this has implications on what we consider “infection” within the seroepidemiological contexts. Our findings may provide a strategy for studies that are focused on discriminating between infection and vaccination using serological endpoints.

## Methods

### Plasmids and Viruses

Recombinant viruses were synthesized using the reverse genetics (rg) method(Hoffmann et al., 2000). Viruses were propagated in embryonated chicken eggs and their sequences confirmed prior to further studies. Viruses were titered by plaque assay on MDCK cells. For immunization experiments, recombinant H3_V_N1 and H6N2_V_ rg viruses bearing either variant H3/constant N1 (H3vN1) or constant H6/variant N2s (H6N2v) representing distinct HA or NA antigenic clusters (Table 1). Four variant H3 or N2 were selected to represent sufficient antigenic distance based on published reports(Bedford et al., 2014; Sandbulte et al., 2011; Smith et al., 2004).

For evaluation of the HA antibody landscapes, a panel of rg H3N2 IAVs bearing the HA and NA from 24 human H3N2 strains from 1968 to 2019 (Supplementary Table 1) were rescued and used as antigens in HI assays. A separate panel of viruses expressing only the human N2 with and the seven genes of A/Puerto Rico/8/1934 (H1N1) (PR8) were rescued for use as antigens in enzyme-linked lectin assay (ELLA).

For virus inactivation, viruses were inactivated with 0.1% beta-propiolactone and concentrated by sucrose density gradient centrifugation. H3 and N2 content were determined by Western blot using anti-H3N2 HA and anti-H3N2 NA antibodies with signal normalized to known concentrations of a standard using HA from A/California/07/2009 (H1N1).

### Growth kinetics

Viral growth kinetics were performed using Madin–Darby Canine Kidney (MDCK) cells and humanized MDCK (hMDCK) cells(Takada et al., 2019). Cells were inoculated with H3_V_N1, H6N2_V_ or PR8 at a multiplicity of infections (MOI) of 0.001. After one hour incubation at 37°C, viral inocula were replaced by viral infection medium (0.3% bovine serum albumin and 1μg/ml TPCK-trypsin in minimal essential medium). Supernatants were collected at 6, 12, 24, 36, 48 and 72 hours post infection (HPI) and viral titers measured by tissue culture infectious dose 50% (TCID_50_) in MDCK cells (Supplemental Fig. 1).

### Hemagglutination inhibition assay

Mouse serum samples were treated with receptor-destroying enzyme for 18 hours and heat-inactivated at 56°C for 30 min. Serum samples were serially diluted two-fold with a starting dilution of 1:10 and incubated for 30 minutes with four agglutinating doses (4AD) of test antigens. 0.75% guinea pig red blood cells in U-bottom 96-well plates according to World Health Organization (WHO) guidelines(Anonymous).

### Enzyme-linked lectin assay

Neuraminidase inhibition (NI) sera antibody titers were determined by enzyme-linked lectin assay (ELLA) as described(Couzens et al., 2014). In brief, 96 well plates were coated with fetuin at 4°C overnight. Mouse sera were heat inactivated and serially diluted from 1:10 to 1:5120 and mixture of virus and sera was incubated for 18 hours in fetuin-coated plates. Plates were washed with 3 times PBST and incubated with PNA-HRP for 2 hours. The NI titers were measured from the serum dilution that yielded 50% inhibition of the virus control.

### HA or NA imprinted mouse models

Six to eight-week-old female BALB/c mice were purchased from Charles River Company (Beijing, China). All animal experiments were performed according to protocols approved by the First Affiliated Hospital of Guangzhou Medical University Institutional Animal Welfare and Ethics Committee.

For sequential priming to induce diverse HA imprinting (**S**equential **H**A, SH), n=10 mice were intranasally (IN) inoculated with 10^5^ PFU of recombinant virus, H3_EN72_N1 and H3_WU95_N1 in 30μL total volume. At 28DPI these mice were IN inoculated with 10^5^ PFU of H3_PE09_N1 and H3_HK14_N1 in 30μL total volume (Supplemental Fig. 2 and 3). Serum samples were collected 28 days after both the first and second inoculations. For control, sequential priming using wild-type like H3N2 viruses (**S**equential **H**A **N**ormal, SHN), n=5 mice were IN inoculated with 10^5^ PFU of H3N2/EN72 and H3N2/WU95 in 30μL total volume. At 28DPI mice were IN inoculated with 10^5^ PFU of H3N2/PE09 and H3N2/HK14 in 30μL total volume. Serum samples were collected 28 days after both the first and second inoculations. The dose of inocula administered to SH and SHN groups were selected based on a preliminary experiment showing that these doses induced comparable antibody responses (data not shown). Further, the limited volume that IN inoculation necessitates limited each inoculation to two different viruses.

For **C**ocktail priming to induce **H**A imprinting (CH) mice were divided into the following groups (n=10 for each group) and inoculated intraperitoneally (IP) with the following: CH1; 10μg of inactivated H3_V_N1 IAV. CH2; 10μg of inactivated H3_V_N1 IAV with Addavax, the squalene-oil-in-water adjuvant, at a 1:1 (v/v) ratio. CH3; H3_V_N1 live viruses at 10^5^ plaque forming units (PFU) each. PBS; inoculated with PBS. A booster dose identical to these initial doses was administered to CH1, CH2 and CH3 28 days post inoculation (DPI). PBS mice were inoculated with PBS at 28DPI (Table 2).

**Table 2.** Study design to generate HA imprinted mice.

|  | **Group** | **DAY 0** | **DAY 28** | **DAY 56** |
| --- | --- | --- | --- | --- |
| **Cocktail priming** | CH1 | inactivated H3<sub>EN72</sub>N1, H3<sub>WU95</sub>N1, H3<sub>PE09</sub>N1, H3<sub>HK14</sub>N1viruses | inactivated H3<sub>EN72</sub>N1, H3<sub>WU95</sub>N1, H3<sub>PE09</sub>N1, H3<sub>HK14</sub>N1viruses | **Challenged with H3N2/SG16 or H3N2/AI68** |
|  | CH2 | inactivated H3<sub>EN72</sub>N1, H3<sub>WU95</sub>N1, H3<sub>PE09</sub>N1, H3<sub>HK14</sub>N1viruses, with adjuvant | inactivated H3<sub>EN72</sub>N1, H3<sub>WU95</sub>N1, H3<sub>PE09</sub>N1, H3<sub>HK14</sub>N1viruses, with adjuvant |  |
|  | CH3 | live H3<sub>EN72</sub>N1, H3<sub>WU95</sub>N1, H3<sub>PE09</sub>N1, H3<sub>HK14</sub>N1viruses | live H3<sub>EN72</sub>N1, H3<sub>WU95</sub>N1, H3<sub>PE09</sub>N1, H3<sub>HK14</sub>N1viruses |  |
| **Control** | PBS | PBS | PBS |  |

For **C**ocktail priming to generate **N**A imprinted mice (CN), mice were divided into the following groups (n=10 for each group) and inoculated IP with the following: CN1; 10μg inactivated H6N2_V_. CN2; 10μg inactivated H6N2_V_ with Addavax at a 1:1 (v/v) ratio. CN3; a cocktail of H6N2_V_ live IAVs at 10^5^ PFU each. PBS; inoculated with PBS. A booster dose identical to these initial doses was administered 28 DPI (Table 3).

**Table 3.**
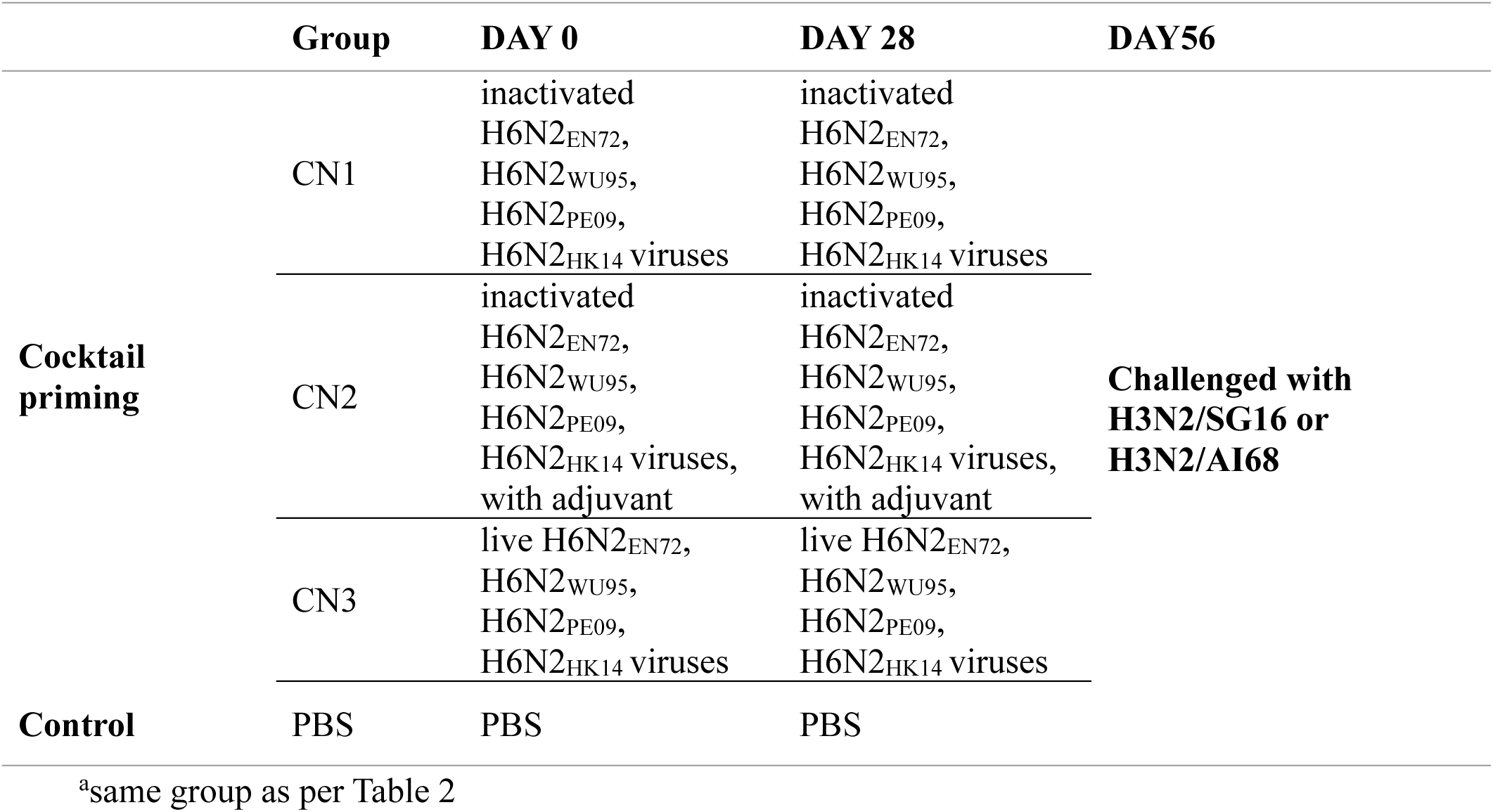
Study design of NA imprinted mice.

Twenty-eight days after final vaccination, all the groups (CH and CN) were challenged intranasally with either rg-derived 10^6^ PFU of A/Singapore/INFIMH-16-0019/2016 (H3N2) (SG16) or 10^4^ PFU of A/Aichi/2/1968 (H3N2) (AI68). Challenge doses were selected as they induced an antibody response and caused detectable weight loss in a preliminary experiment (data not shown). Mice were monitored at least daily and weights were recorded daily for 14 days. Serum samples were collected at the end of the experiment at 28 DPI.

### Computation of antigenic distances in H3 and N2

Antigenic distance (AD) for the tested strains was computed by an amino acid sequence-based approach. The amino acid sequences for H3 and N2 proteins were obtained from the Global Initiative on Sharing All Influenza Data (GISAID, http://gisaid.org) database with their respective accession number (Supplementary Table 2). They were then aligned using MUSCLE(Edgar, 2004), iterated to identify the number of mismatched residues at each antigenic site (Supplementary Table 3) with Python 3. At each antigenic site of H3(Abed et al., 2002; Skowronski et al., 2014), the number of mismatched residues was divided by the total number of amino acids and multiplied by 20 to represent the AD within a 20-dimensional shape space previously described and optimised by Smith et al(Smith et al., 1997). The sum of AD across the five antigenic sites was averaged to compute the AD for the entire antigen. For N2, the antigenic sites were selected from residues reported to be critical to antibody binding(Air et al., 1985; Chen et al., 2018; Colman et al., 1983; Gulati et al., 2002; Kirkpatrick Roubidoux et al., 2021; Lentz et al., 1984; Martinez et al., 1983; Nuss and Air, 1991; Stadlbauer et al., 2019; Wan et al., 2019; Wang et al., 2021; Webster et al., 1982). The sites were not aggregated into groups, so the number of mismatched residues across all antigenic sites was divided by the total number of antigenic sites across the entire antigen, then multiplied by 20.

### Statistical Analysis

Data were analyzed by using GraphPad Prism version 9. HI titers of less than the first dilution of 1:10 were inputted as 5. ELLA antibody titers of less than the first dilution of 1:20 were inputted as 10. All antibody titers were log_10_ transformed. Data were analyzed using unpaired two tailed t-tests.

### Data availability

The datasets generated and analyzed in this study are available on reasonable request.

## Acknowledgements

We gratefully acknowledge Jiaying Jiang and Joyce Chen of Guangzhou Medical University for their technical and administrative assistance. This research was funded by Natural Science Foundation of Guangdong Province, China, Grant No.2019A1515012070 and Guangdong Provincial Research and Development Projects in Key Areas (Grant No.2019B111103001). SW and BC are supported by a grant from the Research Grants Council of the Hong Kong Special Administrative Region, China (Project No. T11-712/19-N).

## Author Contribution

Study conceptualization: SW, XL and MZ; Performed the experiments and collected data: XL, JW, SL, CX, YB, BX; Data analysis: XL; Manuscript preparation: XL, SW, MZ, BC, BY, RW; Final Draft: LZ, RW, SSW, MZ; Funding acquisition: SSW, BC. All authors have read and approved of the final manuscript.

## Supplemental Figures

**Supplementary Fig. 1.**
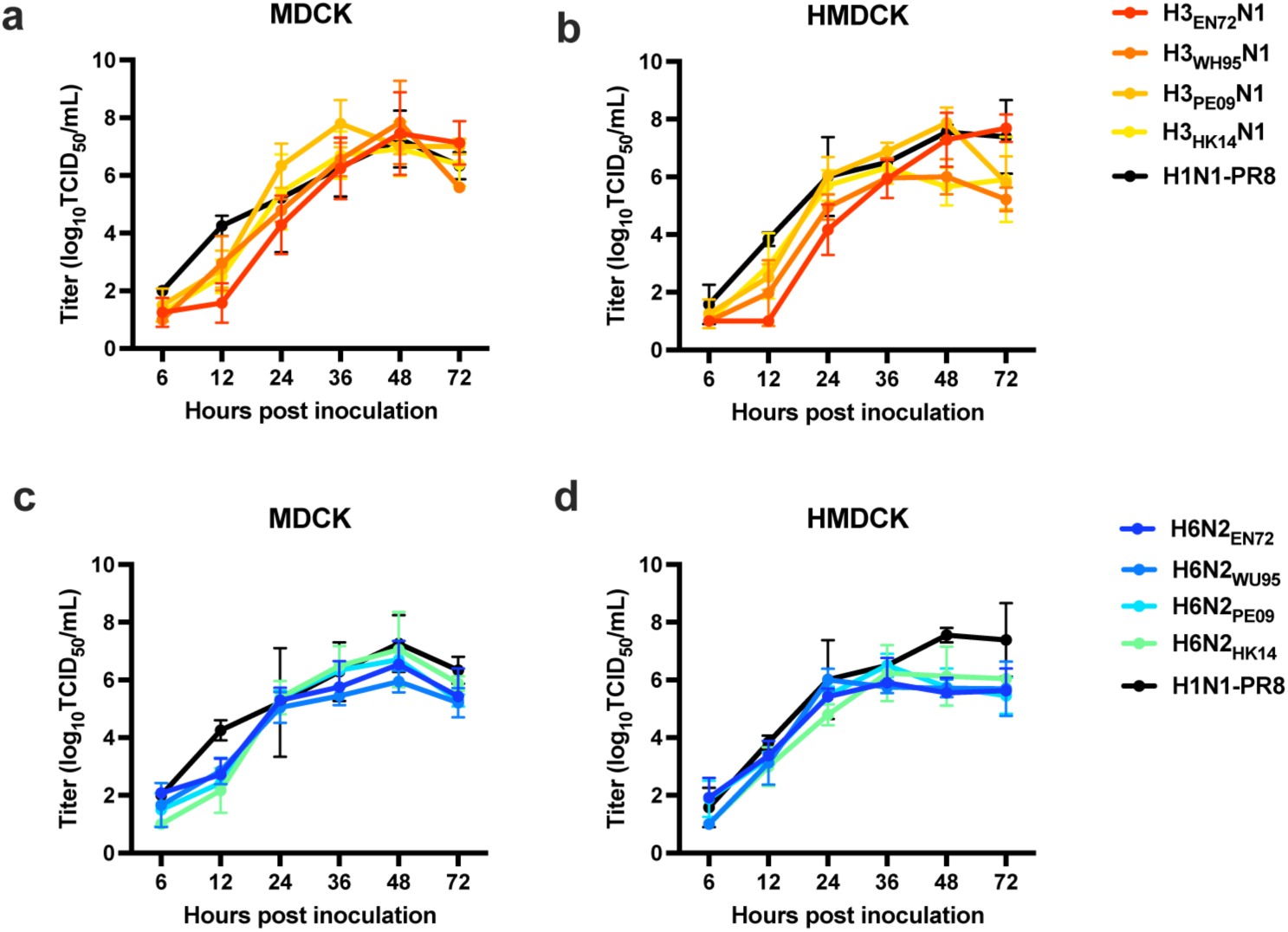
Replication kinetics of H3_V_N1 and H6N2_V_ viruses in MDCK and HMDCK cells. **a-b** MDCK cells and HMDCK cells were infected with H3_EN72_N1, H3_WU95_N1, H3_PE09_N1, H3_HK14_N1 and H1N1-PR8 (MOI=0.001). Supernatants were collected at different time points and determinate by TCID_50_ assay. **c-d** Growth kinetics of H6N2_EN72_, H6N2_WU95_, H6N2_PE09_, H6N2_HK14_ and H1N1-PR8 in MDCK cells and HMDCK cells.

**Supplementary Fig. 2.**
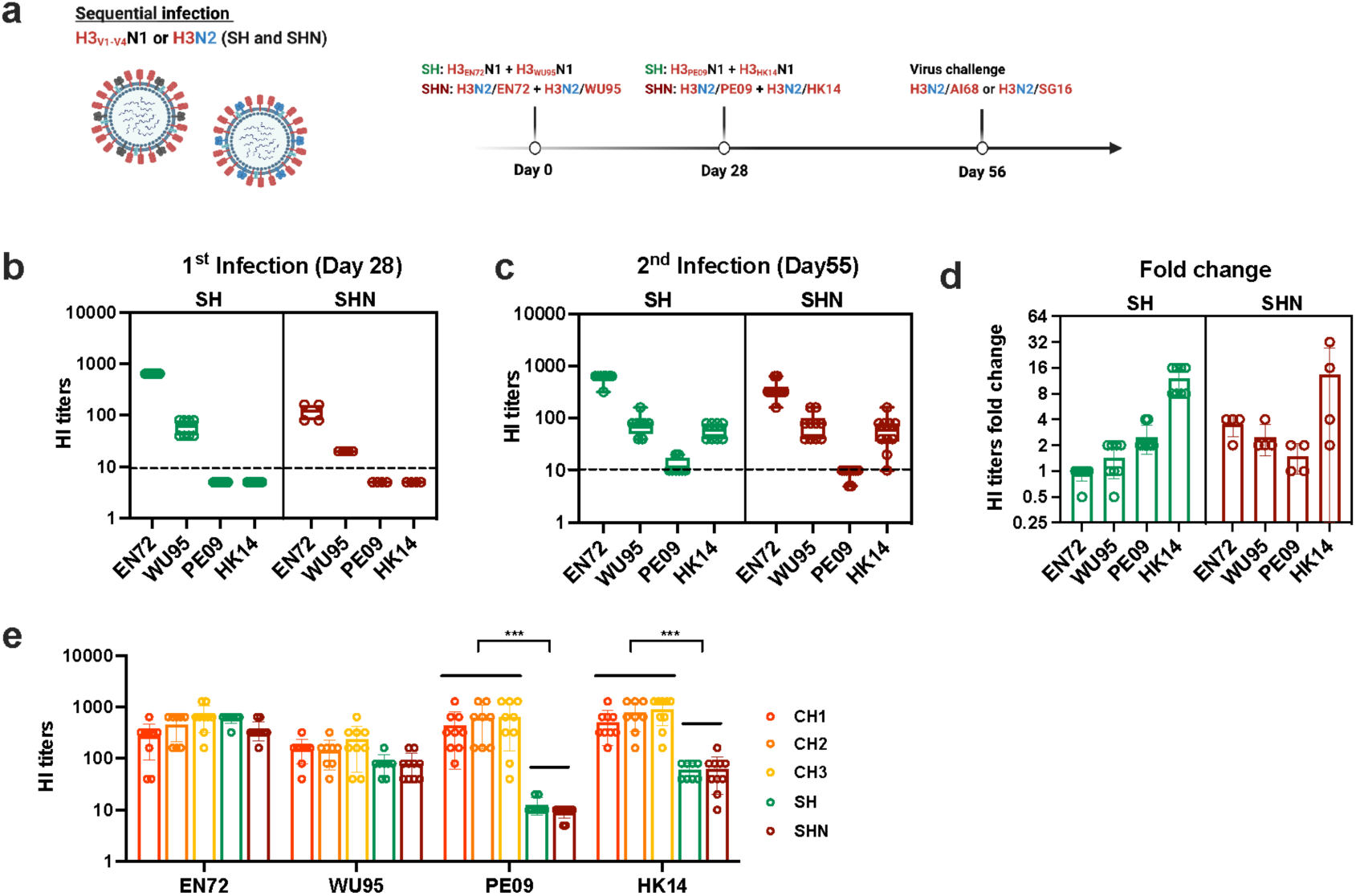
Serum antibody response after sequential priming with diverse H3. **a** Schematic of sequential infection. For the **s**equential **H**A priming (**SH**) group experiment, H3_EN72_N1 and H3_WU95_N1 were inoculated first, intranasally, followed by H3_PE09_N1 and H3_HK14_N1. This two-dose strategy was adopted as the inoculum volume could not accommodate the desired infection dose for the four viruses. As control, we also intranasally inoculated wild-type like (**N**ormal, i.e bearing the original H3 and N2 genes) rg-viruses in the same sequential order (**SHN**). **b-c** Hemagglutination inhibition (HI) serum antibody titers against EN72, WU95, PE09 and HK14 were determined after 1^st^ and 2^nd^ infection. **d** HI antibody fold change between 1^st^ and 2^nd^ infection. **e** HI titers in cocktail and sequential priming mice after 2^nd^ dose or infection. * indicates *P*-value <0.05, ** indicates *P*-value <0.01 and *** indicates *P*- value <0.001 by using unpaired two tailed t-tests. Dotted lines indicated HI titers of 1:10.

**Supplementary Fig. 3.**
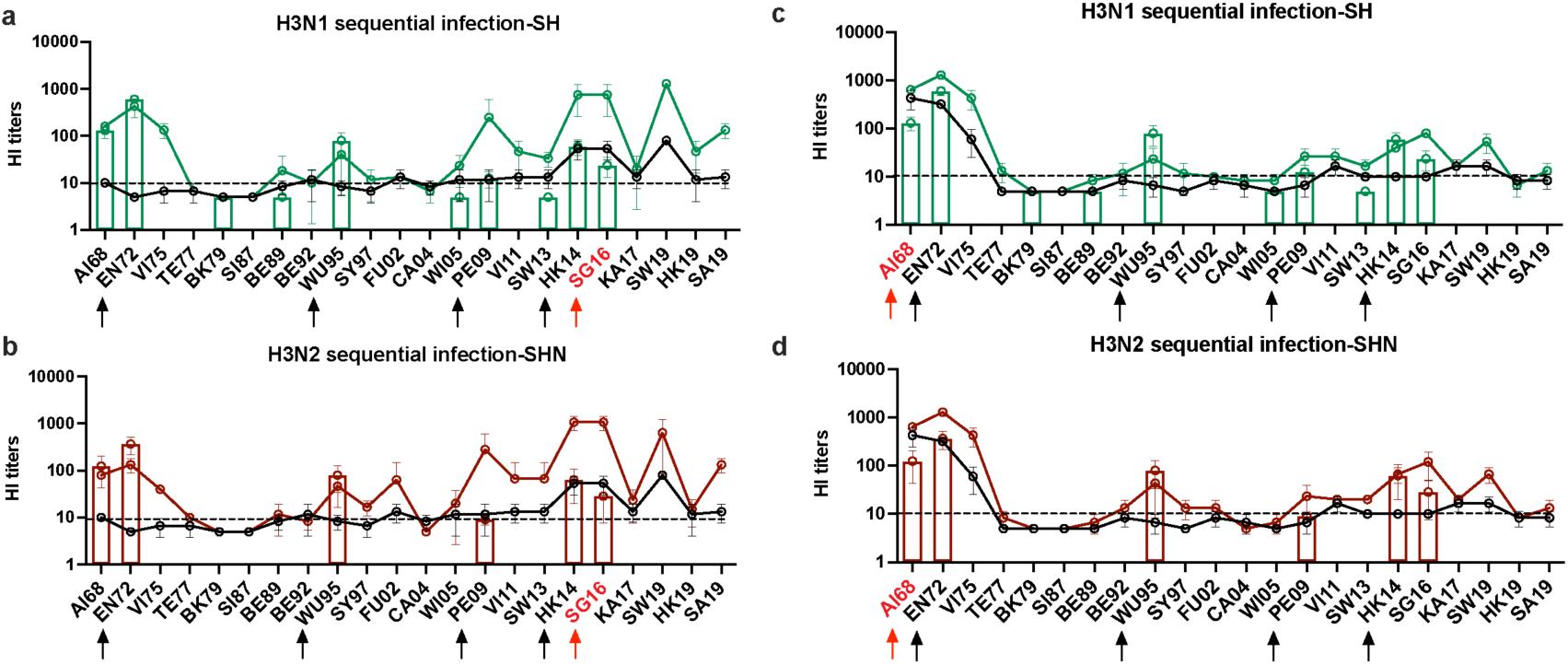
HI landscape after infected with SG16 or AI68 in sequential priming mice. 21 days after final infection, HI antibody landscape against 22 historical H3N2 viruses was detected in sequential priming mice after infected with 10^6^ PFU SG16 (a-b) and 10^4^ PFU AI68 (c-d). The black line indicated titers in the PBS group challenged with H3N2/SG16 or H3N2/AI68. The red arrow indicated challenge virus. Dotted lines indicated the 1:10 in HI titers.

**Supplementary Table 1.** H3N2 strains used in this study to generate the HA and NA antibody landscape.

| Clade | Strain Name | Abbreviation |
| --- | --- | --- |
| 3C.2a4 | A/Aichi/2/1968 | AI68 |
| 3C.3a | A/England/42/1972 | EN72 |
| 3C.3a | A/Victoria/3/1975 | VI75 |
| 3C.3a | A/Texas/1/1977 | TE77 |
| 3C.3a | A/Bangkok/1/1979 | BK79 |
| 3C.2 | A/Sichuan/2/1987 | SI87 |
| 3C.3a | A/Beijing/352/1989 | BE89 |
| 3C.3a | A/Beijing/32/1992 | BE92 |
| 3C.3a | A/Wuhan/359/1995 | WU95 |
| 3C.2 | A/Sydney/5/1997 | SY97 |
| 3C.2 | A/Fujian/411/2002 | FU02 |
| 3C.3a | A/California/7/2004 | CA04 |
| 3C.3a | A/Wisconsin/67/2005 | WI05 |
| 3C.2 | A/Hong Kong/218849/2006 | HK06 |
| 3C.3a | A/Brisbane/10/2007 | BR07 |
| 3C.3a | A/Perth/16/2009 | PE09 |
| 3C.2 | A/Victoria/361/2011 | VI11 |
| 3C.3a | A/Switzerland/9715293/2013 | SW13 |
| 3C.2 | A/Hong Kong/4801/2014 | HK14 |
| 3C.2a.1 | A/Singapore/INFIMH-16-0019/2016 | SG16 |
| 3C.3a.1 | A/Kansas/14/2017 | KA17 |
| 3C.3a.1 | A/Switzerland/8060/2017 | SW19 |
| 3C.2a1b.1b | A/Hong Kong/2671/2019 | HK19 |
| 3C.2a1b.2b | A/South Australia/34/2019 | SA19 |

**Supplementary Table 2.** Accession number for the amino acid sequences of HA and NA proteins used to compute the antigenic distances for tested strains.

| <u>Strain</u> | <u>HA sequence accession number</u> | <u>NA sequence accession number</u> |
| --- | --- | --- |
| AI68 | EPI381955 | EPI381957 |
| EN72 | EPI381987 | EPI381989 |
| BK79 | EPI367158 | EPI367160 |
| BE89 | EPI362508 | EPI362510 |
| WU95 | EPI397298 | EPI362822 |
| WI05 | EPI160218 | EPI160220 |
| PE09 | EPI182941 | EPI182942 |
| SW13 | EPI696955 | EPI696961 |
| HK14 | EPI741474 | EPI741475 |
| SG16 | EPI780183 | EPI810155 |

**Supplementary Table 3.** Antigenic sites of H345 (A) and N247-58 (B)

| <u>Protein</u> | <u>Antigenic site</u> | <u>Residue (in H3 numbering)</u> |
| --- | --- | --- |
| H3 | A (n = 19) | 122, 124, 126, 130-133, 135, 137, 138, 140, 142-146, 150, 152, 168 |
|  | B (n = 22) | 128, 129, 155-160, 163-165, 186, 190, 192-194, 196-198 |
|  | C (n = 27) | 44-48, 50, 51, 53, 54, 273, 275, 276, 278-280, 294, 297, 299, 300, 304, 305, 307-312 |
|  | D (n = 41) | 96, 102, 103, 117, 121, 167, 170-177, 179, 182, 201, 203, 207-209, 212-219, 226-230, 238, 240, 242, 244, 246-248 |
|  | E (n = 22) | 57, 59, 62, 63, 67, 75, 78, 80-83, 86-88, 91, 92, 94, 109, 260-262, 265 |

B:
| <u>Protein</u> | <u>Residue (in N2 numbering)</u> |
| --- | --- |
| N2 | 118, 119, 125, 150-152, 197-199, 220-222, 224, 245-249, 253, 258, 272, 277, 295, 296, 313, 329, 331, 334, 339-347, 356, 368, 369, 370, 371, 400, 403, 406, 429, 431-434, 468 |

